# Electrical diversity of neurons in sensory cortices

**DOI:** 10.1101/2024.10.07.617007

**Authors:** Niccolò Calcini, Fleur Zeldenrust, Tansu Celikel

## Abstract

The classification of neuronal types is a complex task with numerous molecular, anatomical, and functional (electrical) features have been identified as informative in discriminating neuronal populations. The functional characterization of neurons has traditionally been carried out with predefined sets of parameters such as firing rate and action potential generation threshold. Here we provide an objective method to choose what parameters are most informative about a neuron’s functional cell type. Using this method we show that despite the significant molecular and anatomical variability across neurons, functional characterization of neuronal activity identifies 9 and 11 distinct neuronal subpopulations in the upper layers of the somatosensory and visual cortices respectively. This novel classification method will help to unravel the functional (electrophysiological) diversity of cellular classes throughout the nervous system. Further, the thorough comparison between the different classes of cells will provide a solid building block for the study of the sensory cortices.

## Introduction

The neurons in layers (L) 2/3 of the barrel cortex of rodents are involved in integrating information that originates from the sensory periphery with the information generated elsewhere in the brain (1; 2; 3; 4). They carry spatial and temporal information about contacts with an object, tactile features of the object and the whisker position in parallel (5). Equivalently, neurons in the supragranular layers (i.e. L2/3) of the primary visual cortex (V1) integrate and carry information about visual features such as orientation, movement, density and contrast (6; 7; 8). Given their importance of the single building blocks of the sensory circuits for the processing of sensory information, one of the major goals of systems neuroscience is to describe and classify them.

Neurons are commonly classified according to their anatomical features, molecular fingerprints, as well as passive and active electrical properties (9; 10; 11; 12; 13; 14; 15; 16; 17). The considerable molecular and biochemical diversity of neurons (9; 10; 14; 16; 18) necessitates multi factorial classification of neurons based on molecular fingerprints and anatomical features which however does not necessarily correspond to the same variety in functional properties. For example, neurogliaform neurons can be classified as a single subpopulation of inhibitory neurons when both their anatomical features and the expression of the 5-hydroxytryptamine receptor 3A receptors are considered. However, because these neurons also express Glutamate Decarboxylase 1 and Vasoactive Intestinal Peptide at varying rates, the neurogliaform neurons can be further sub-classified (14). Furthermore, the expression of the molecular markers is often subject to experience-dependent plasticity, therefore, cellular classification based solely on molecular markers might confound the clustering efforts. Considering that action potential rate and timing are the means of neuronal communication in synaptically coupled networks, active electrical properties of neurons could instead be used to determine the functional subclasses of neurons.

Cellular classification based on active electrical properties of neurons necessarily requires redefining experimentally recorded continuous voltage (or current) time-series into variables (also known as descriptors, features, dimensions), e.g. action potential threshold, amplitude, spike half-width, usually via their mean or median values. This approach, combined with various clustering methods, has been used to electrically classify neurons before, although with arbitrarily selected features, or with features with low interprepatibilty (10; 19; 20; 21; 22; 23; 24). Here we propose an objective method of feature selection, identify a subset of physiologically interpretable features after dimensionality reduction and subsequently rank these features according to their performance in defining the functional clusters.

In previous clustering studies, the neural responses to sustained somatic activation (or inactivation) have been studied upon depolarization (or hyperpolarization) due to a constant stimulus intensity. However, given the prominent voltage-gated conductances the integrative properties of neurons are likely to vary depending on the stimulus amplitude. For this reason, and not to limit bias the feature selection to a specific level of current injection, we included as clustering variables, also the adaptive changes in spiking features upon systematic changes in somatic depolarizations. Clustering performed after the identification of the features that describe supra- and subthreshold voltage dynamics within and across different depolarizing stimulus intensities showed that adaptive changes in voltage dynamics are a powerful predictor of neuronal classes both in somatosensory and primary visual cortices. The results of the classifications performed on these datasets showed that the supragranular layers contain ≃ 11 electrically distinguishable neuron classes independent from the sensory cortex studied.

## Methods and Materials

All data analysis were performed in MATLAB using custom-written routines (https://github.com/DepartmentofNeurophysiology/ Voltage traces corresponding to somatic current injection epochs (500 ms/sweep) were used for the quantification of the active electrophysiological properties of neurons (Figure 1)

**Figure 1.**
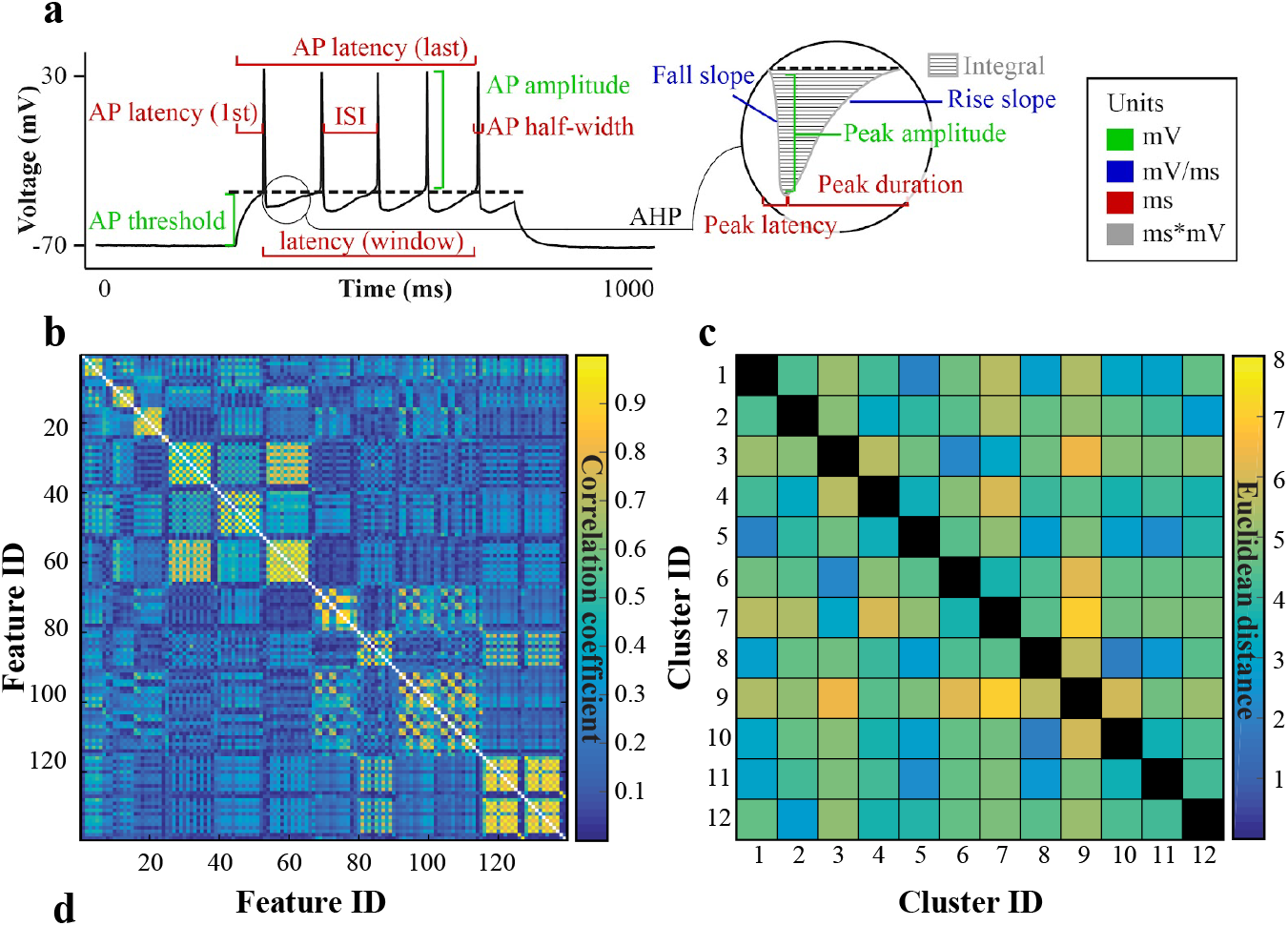
Feature space, dimensionality reduction. **A)** An example voltage trace and the description of selected primary features. AP = action potential, ISI = interspike interval; AHP = afterhyperpolarization peak. **B)** The pairwise Pearson correlation coefficient for 139 variables represented as a matrix (absolute value scale). **C)** The pairwise Euclidean distance between the cluster centroids after standardization in the S1 dataset.

### Data description

The “S1 dataset” (N = 220 neurons) was made freely available by our group before (da Silva Lantyer et al., 2018). The data consists of whole cell current-clamp recordings from acute brain slices coming from Pval-cre and SSt-cre mice from both sexes, aged between 9 and 54 weeks (see (da Silva Lantyer et al., 2018) for raw data, experimental details and basic analysis routines). The V1 dataset (N = 282 neurons) was collected and distributed by the Allen Brain Institute. It is available for download at http://celltypes.brain-map.org/. The associated experimental procedures are available online as a white paper (http://celltypes.brain-map.org/#ephys, last accessed on 14/03/2021). We have isolated experiments from adult mice (Mus musculus, different Cre-driver lines), aged between 4 and 10 weeks. Only recordings from the primary visual cortex (V1) were included in the analysis. From this “V1 database”, we used the current-clamp protocol named “Long square”, where 1 s or 2 s long square (step and hold) current pulses were injected to depolarize the somatic membrane potential, which was otherwise clamped at -70 mV. To allow for a direct comparison between the two datasets, we analyzed only the first 500 ms of the V1 square pulse, as that was the length of the step and hold stimulus in the S1 dataset. The number and amplitude of the current injections were not constant across different experiments. To ensure that a similar analysis can be employed in both datasets, we binned the data into current injections steps of 40 ± 15 pA, between +40 and +400 pA. If one more than one stimulus conditions fell within a given bin, only the smallest current injection was included.

### Feature space

All data analyses were performed in MATLAB using custom-written routines available at (https://github.com/DepartmentofNeurophysiology/NClust). Voltage traces corresponding to somatic current injection epochs (500 ms/sweep) were used for the quantification of the active electrophysiological properties of neurons (Figure 1A, Supplementary Table 1). A total of 139 features/neuron were extracted. Seventy of these features were descriptors of the sub- and suprathreshold responses measurable in a single stimulus presentation. The remaining 69 captured the dynamical nature of the change in somatic voltage dynamics across multiple depolarization states: to represent this, we used a linear fit of the features measured over the different current intensities used in the experimental protocols. The linear fit was used after a “best fit” analysis. The results showed that the overwhelming majority (i.e. 67/70) of the dynamic changes in the features could be best approximated by a linear fit with varying slopes. To ensure a uniform comparison across the feature space, we fitted all dynamical parameters (i.e. varying with stimulus amplitude) with a linear kernel, and used the slope and onset value of each fit as descriptors, obtaining the total of 139 parameters (the parameter representing the “lowest current step at which the cell fired an AP” does not have a fit value).

**Table 1.**
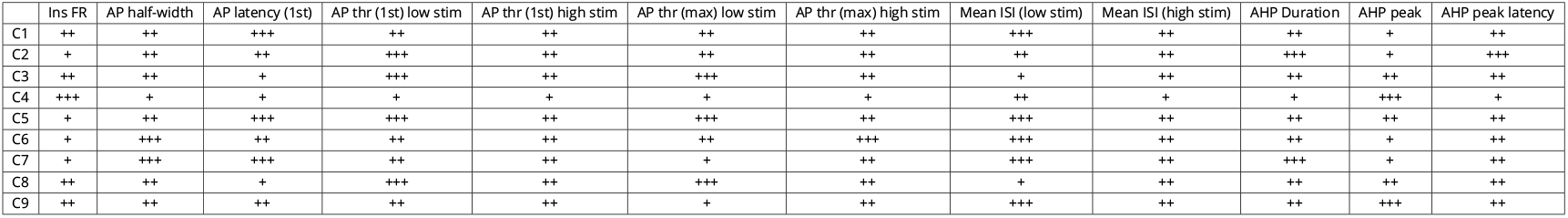
Clusters characteristics described as 3-bin values. The ‘+’ represent: for Instantaneous FR and AP-half-width: low (+), mid (++), high (+++); for AP and AHP latency: + (short latency), ++ (mid), +++ (slow=longest latency); for AP threshold: + (hyperpolarized), ++ (mid), +++ (depolarized); for ISI and AHP duration: + (short), ++ (mid), +++ (long); for AHP peak: + (small), ++ (mid), +++ (large). Detailed description of the clusters features across different stimulus levels are shown in Supplementary Figure 2. S1 clusters.

### Dimensionality reduction

To estimate the minimum number of dimensions required to describe the data, we performed a Principal Component Analysis (PCA). We computed the eigenvalues of the whitened PCA covariance matrix, obtaining the lower boundary estimation, i.e. 15, as described before in Gao et al. 2017 (25):

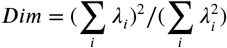

To reduce the dimensionality in the feature space, we applied empirically defined thresholds on the Pearson correlation coefficients computed pairwise across all parameters, and removed all the redundant parameters which displayed significant pairwise correlation. 40 and 48 parameters were selected with this method for the S1 and V1 datasets, respectively (see Figures 1, 2). Given the lower limit estimate of the dimensionality of ∼ 15 (∼ 14.5 and ∼ 15.5 for S1 and V1, respectively) and datasets of 220 and 282 neurons, this feature space represents a sufficiently small dimensionality to perform the initial clustering.

**Figure 2.**
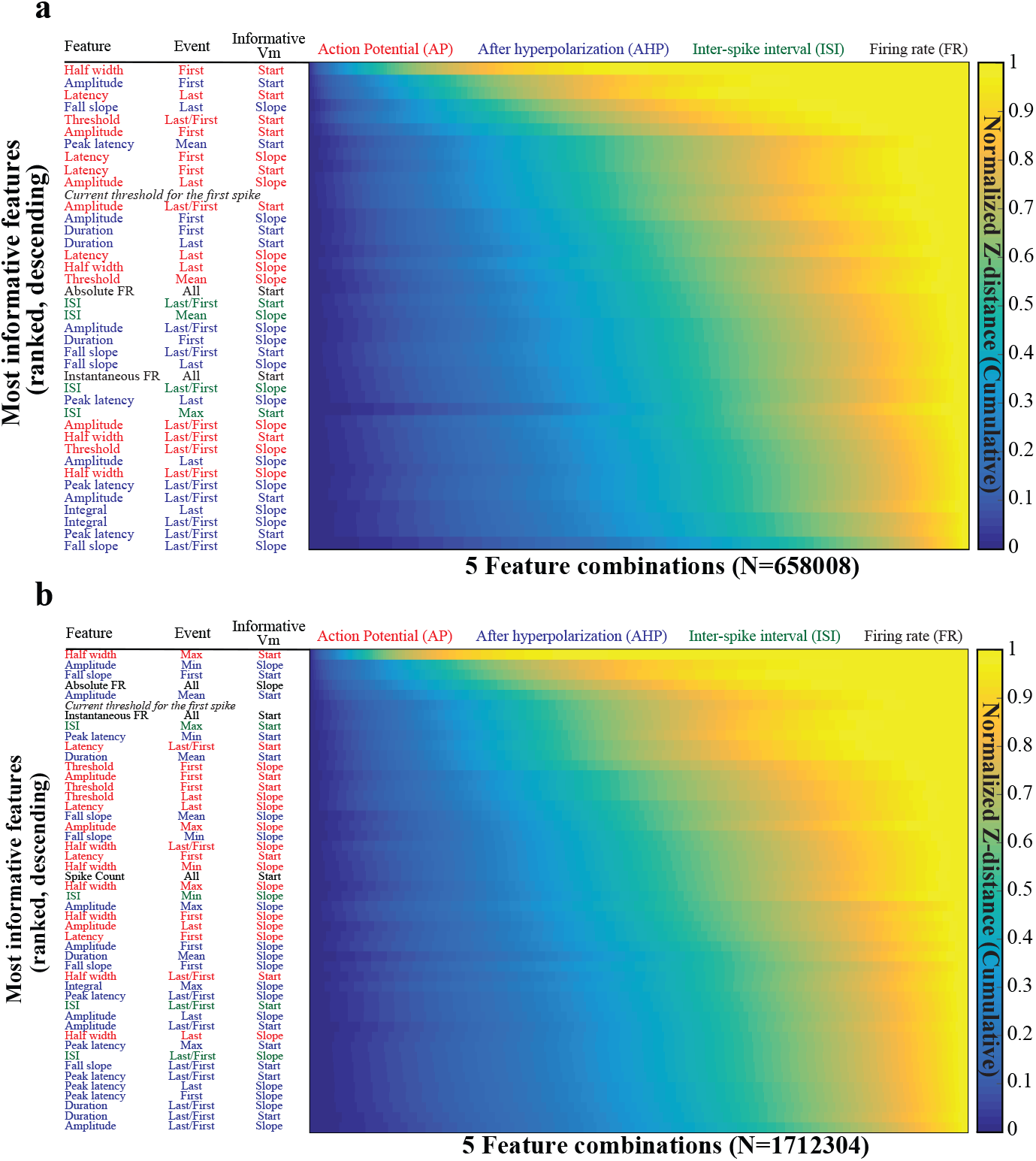
Features ranking. **A)** Ranking of the features surviving the correlation based dimensionality reduction, for the S1 dataset. Each row represents the cumulative sum of that feature count, in the 5 parameters combinatorial space ordered by mean Z-distance between clusters, so that a feature appearing earlier would scale higher. **B)** Ranking of the features for the V1 dataset

### Feature ranking

To systematically evaluate what features contain most information for the electrical classification of neurons, and provide a simplified set of criteria for post-hoc replicability across different datasets, we computed the average Z-distance between any given clusters across any combination of 5 features that survived the dimensionality reduction described above. As the final step, we ran the standard k-means algorithm, with the number of clusters estimated with the maximum likelihood agglomerative clustering for all feature sets (S1: N = [139 40 5], V1: Ns = [139 48 5]) and compared them by computing the pairwise Z-distance between any given two clusters (Figure 1 B, C). To quantify the clustering performance with a subset of features, we repeated the clustering one final time but with the combination of five features that allowed for maximal cluster separation (i.e. slope of mean inter spike interval (ISI), onset values of ratio between threshold on first and last APs, latency of last AP, half-width of 1st AP and AHP peak amplitude, for the S1 database. Slope of absolute firing rate and AHP minimum peak amplitude, onset values of the maximum AP half-width, mean AHP peak amplitude, 1st AHP fall slope, for the V1 database.). The results showed that all neurons had a unique cluster identity, suggesting that in a five-dimensional space neurons can still be grouped with high-confidence into well-separated clusters (Figure 2).

### Clustering

To sort neurons into cell-classes, we used a hierarchical clustering approach where neurons were first sorted into a minimum of 50 classes using k-means clustering (45). Note that a subset of the cells could not be clustered due to missing observations in their 139 dimensional description, for example because of lack of spiking during small amplitude current injections. They were subsequently merged iteratively using maximum likelihood estimations as described below and before (46). K-means clustering was performed after standardizing the data so that each feature had a mean of zero and a standard deviation of one. After clustering, those cell-clusters that had less than 10 cells were removed and its members were reassigned to the closest cluster based on Z-distance minimization (25; 26). This process was repeated 10000 times to calculate the variance in cluster assignments, which was subsequently used to determine the final cell-clusters based on maximum likelihood (ML) estimations, assigning each cell to the cluster with the highest probability of containing it. If, after the ML assignment, there were any clusters with less than 10 cells, the entire procedure was repeated again but this time ML clusters were used as the starting clusters.

## Results

To estimate the number of cell classes in the supragranular layers of the adult mouse barrel cortex (S1) and primary visual cortex (V1), we employed a hierarchical clustering approach on voltage traces recorded intracellularly in acute cortical slices. We ranked each electrophysiological parameters based on their performance in defining each cluster (distance between clusters), and show how it is possible to achieve an efficient (functional) clustering of both datasets based on a the combination of 5 paramters which allows for the best cluster separation. We characterize and compare the clusters obtained in both the S1 and V1 datasets, and, finally, showed how the clustering can, in fact, be greatly affected by the stimulus intensity.

### Dimensionality reduction for neuronal classification

The neuronal response to somatic current injections was approximated by 139 dimensions that represent the response dynamics within and across stimulus intensities (i.e. the amplitude of the current injection). To determine whether neurons can be clustered with high confidence in this high-dimensional feature space, we performed k-means clustering and ML-based cluster merging (see Materials and Methods for details). The quantification of the occupancy for each cluster (Supplementary Figure 1A) showed a heteroscedastic distribution. The probability of a given neuron to be assigned to any given cluster (Supplementary Figure 1B) did not follow a uniform distribution, and only a subset of neurons was clustered with high confidence (Supplementary Figure 1C). These observations support the notion that clustering in a high-dimensional space results in clusters with a small (minimal) inter-cluster distance. Accordingly, under these constraints, cluster identification is a probabilistic inference where each neuron has a particular probability distribution to be associated with a finite number of clusters (Supplementary Figure 1C).

The clustering performance can be improved by either reducing the dimensionality of the feature space or by increasing the sampling size (i.e. number of neurons) to minimize the distance between the members of a given cluster while increasing the inter-cluster distance. Given the 220 (S1) and 282 (V1) neurons available in the two datasets, we opted to reduce the dimensionality of the feature space.

To estimate the minimum number of features required to describe the data, we computed the dimensionality of the dataset based on the eigenvalues of the covariance matrix (see Materials and Methods for details), which showed that a minimum of 15 features is required to maximally reconstruct the data defined in 139 dimensions. To identify these features we computed the pairwise Pearson correlation across all variables. The correlation matrix (Figure 1B) showed that 12 subsets of features are highly correlated in both datasets. Using an empirically determined threshold of 0.55, we identified “weakly” correlated parameters (N=40 in S1 and N=48 in V1) and repeated the neuronal clustering in this lower-dimensional feature space. For this intermediate step, the 220 neurons from the S1 database were grouped into 12 clusters (Figure 1C) that are well separated from one another (Figure 1C). We obtained a similar result of 11 clusters in the V1 database of 282 neurons.

The 40 (S1) and 48 (V1) features used for clustering are still significantly larger than the theoretical minimum number of features (i.e. 15) required to describe these datasets. To rank the relevant contribution of each feature to the clustering performance we repeated the clustering, but using only 5 features at a time, in a combinatorial space. Across the 658008 and 1712304 combinations in the S1 and V1 datasets respectively, all informative features are the variables which approximate the cell behavior across current injection steps (Figure 2A, B).

### Classification of adult supragranular layer neurons in sensory cortices

To identify the functional classes of adult cortical neurons in the supragranular layers of the primary somatosensory cortex, we deployed the hierarchical clustering described above and assigned each neuron to a cluster. Nine clusters contained 10 or more units. Applying the same method to the V1 dataset, we find a similar result of 11 clusters containing 10 or more units. To systematically compare the changes in the spiking responses across the membrane states, we described the electrical responses to somatic current injections using features that are among the most informative dimensions for clustering (Figure 2).

As previously shown (10; 22; 23; 27), binary classification of putative excitatory (regularly spiking) and putative inhibitory (fast-spiking) neurons can be achieved using firing rate/inter-spike interval, action potential half-width, or any combination thereof (Table 1, Table 2, Figure 3, Supplementary Figure 2). The latency of the after-hyperpolarization (AHP) peak allows further classification of the putative inhibitory neurons into three sub-classes based on the rate of adaptation upon somatic depolarization (Table 1, Table 2, Figure 3). Although all three sub-classes have a comparable AHP duration, which is shorter than that of all putative excitatory neurons, and quickly reach the peak latency during membrane repolarization, their AHP latency follows a gradient and is not suppressed considerably with the increased depolarization of the membrane (Table 1, Figure 3, Supplementary Figure 2).

**Table 2.**
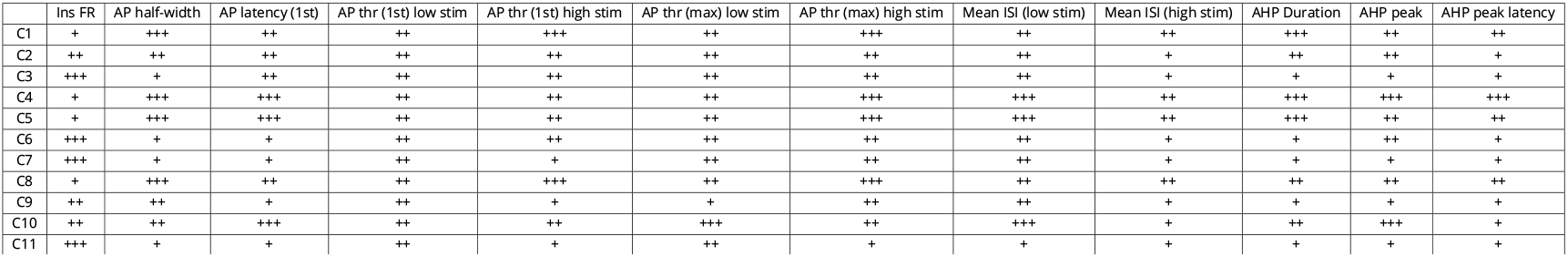
Same as Table 1. V1 clusters.

**Figure 3.**
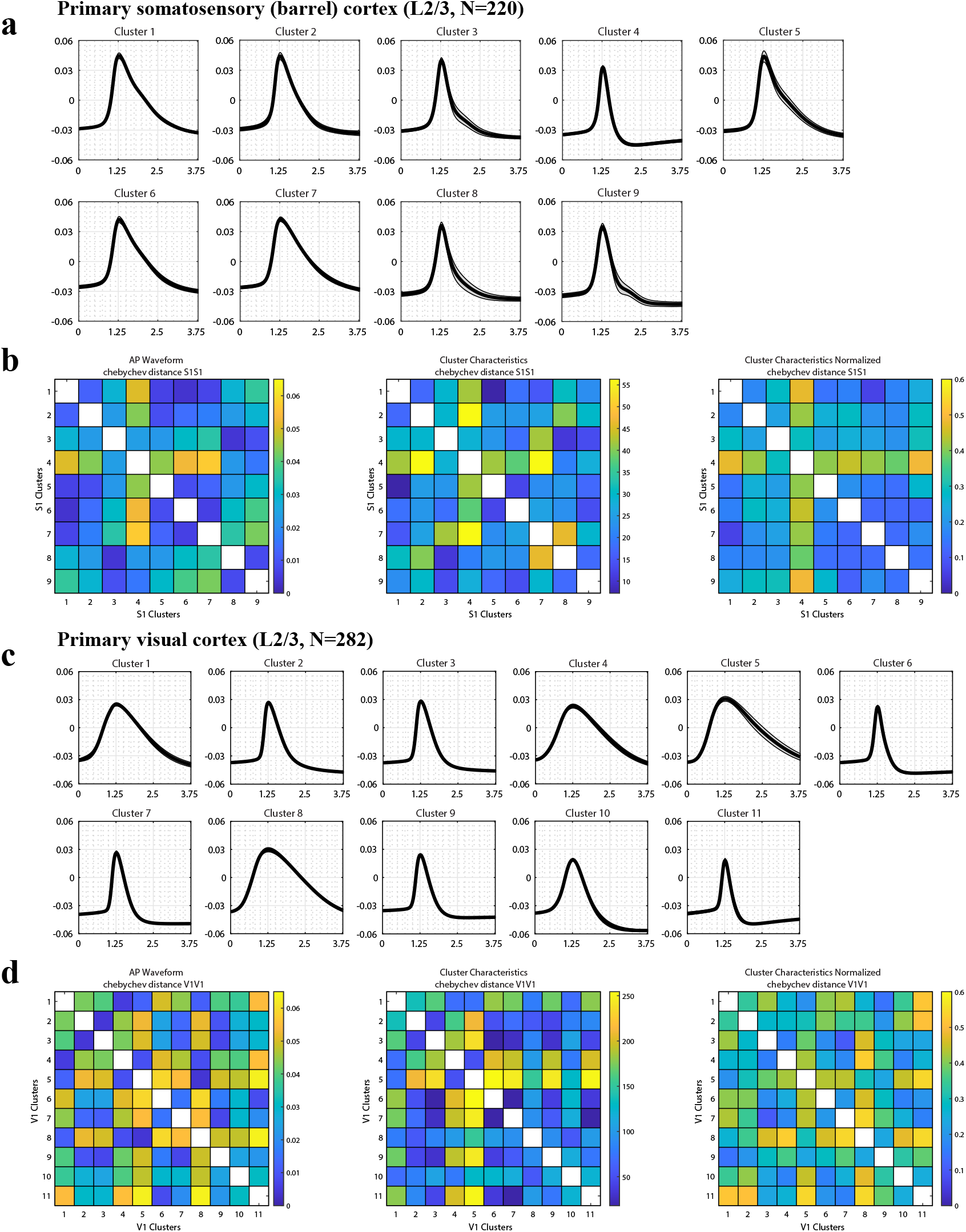
**A)** Average action potential (AP) waveform for the clusters of the somatosensory dataseta. **B)** Chebychev distance between the clusters. From left to right: distance between the AP waveforms; distance between the array of the average features of each cluster (see Table 1, 2); distance between the array of the average, normalized features of each cluster. **C, D)** Same as A, B, for the visual cortex dataset.

**Figure 4.**
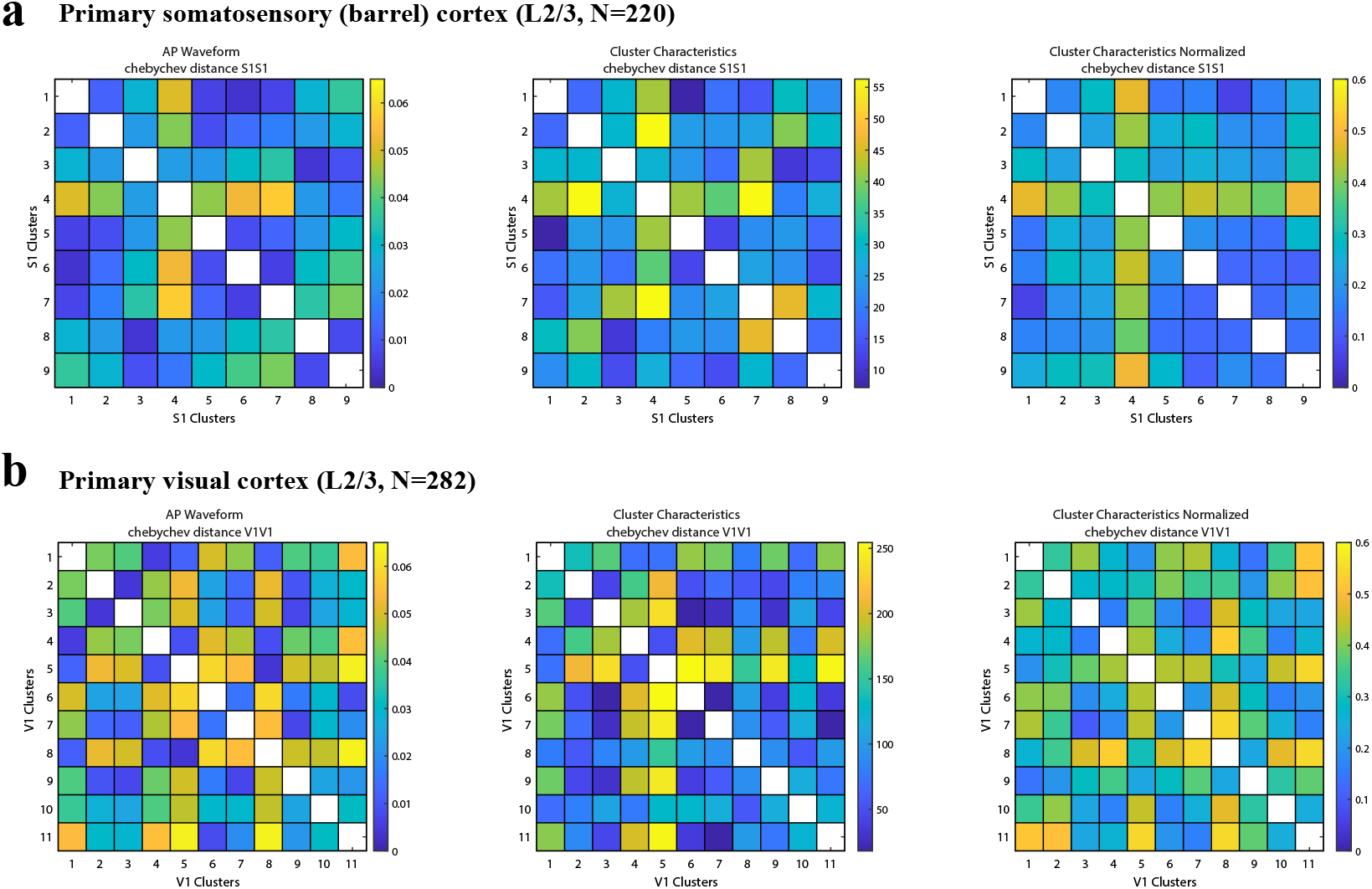
The Chebychev distance between clusters belonging to the same cortical area. The distances were computed between the average AP waveforms of each cluster, and between the vectors containing the means of some of the main features describing the electrophysiological characteristics of the clusters (see Table 1, 2). For comparison, the Chebychev distance was computed also between the same vectors of features with values normalized between 0 and 1.

**Figure 5.**
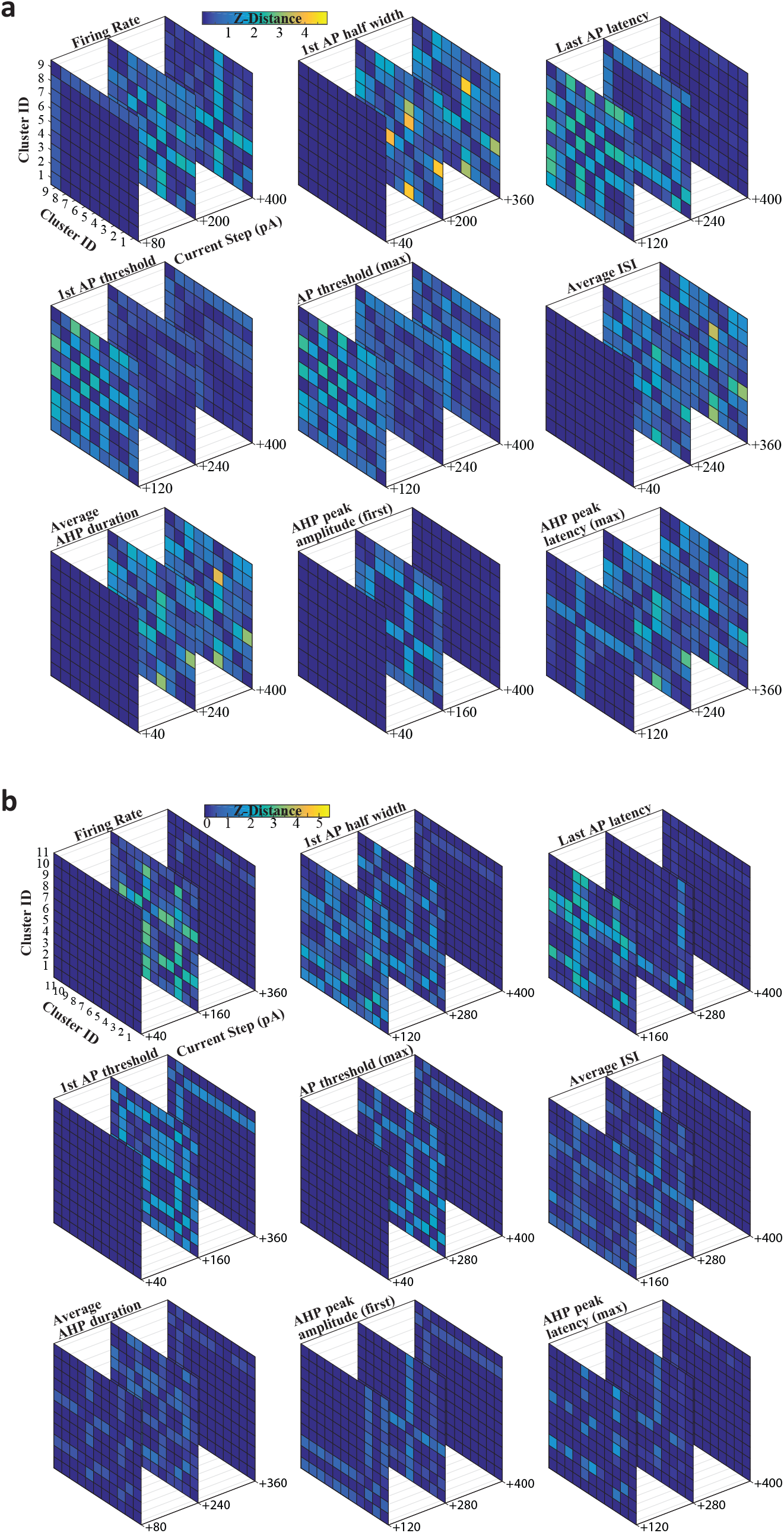
Cluster separation varies with membrane polarization state. The Z-distance between clusters were calculated across all clusters and across stimulus amplitude. Nine representative features and separation distance between clusters are shown. A) Data from the S1 database. B) Data from the V1 database.

In terms of molecular and biochemical complexity, inhibitory neurons are significantly more diverse than their excitatory counterparts (9; 16; 14; 10; 18). Electrophysiologically, however, our findings now indicate that excitatory neurons are at least as diverse as their inhibitory counterparts both in S1 and V1 (Table 1, Figure 3, Supplementary Figure 2). Six of the ten clusters generate action potentials at lower rates than putative inhibitory neurons, and their AP width is broader of inhibitory neurons. The significant variability among excitatory neurons’ spiking patterns (Table 1, Figure 3, Supplementary Figure 2) might be attributed to the properties (i.e. subchannel composition), density and distribution of voltage-gated channels (e.g. in the case of action potential half-width, action potential latency, action potential threshold), the adaptive changes in voltage-gated conductances across depolarization states (resulting in action potential threshold variation) or differences in the membrane integration time constant (e.g. interspike interval). This variability between the putative excitatory neurons might also be due to anatomical or experience-dependent changes across cortical neurons (9; 35). Independent from the underlying cause, these results argue that adult excitatory neurons are as diverse as the inhibitory neurons in the primary somatosensory and visual cortices.

### Neuronal electrical identity depends on the somatic depolarization state

The feature space that best predicts the clustering performance, i.e. most informative feature combination, includes variables that are analytical descriptions of the adaptive changes observed in spiking across somatic (membrane) depolarization states (Figure 1B). This observation suggests that the separation between clusters might depend on stimulus amplitude. Because voltage-gated conductances help maintain membrane depolarization, ensure repolarization and contribute to the action potential generation, sustained depolarization of the membrane potential might change the electrical signature of the neuron (30; 31; 32). Therefore, to quantify the discriminability of the clusters across the discrete somatic depolarization states, we calculated the average pairwise Z-distance between clusters (Figure 3).

## Discussion

The responses of (cortical) neurons are shaped by both their intrinsic biophysical properties and their position and connectivity in the network (33). During development, but also throughout the adult life of an animal, several mechanisms including homeostatic and neural plasticity mechanisms adapt neurons’ physiological properties to the ever-changing cellular and network properties on different timescales (from milliseconds to days), including but not limited to biophysical properties (i.e. what ion channels and receptors are expressed in the membrane and in what quantities) (34), network connectivity (i.e. to what neurons it is connected and how strongly) and cellular morphology. Therefore, we expect that structural and functional correlations between neurons (22; 36) result in neural properties that group together so that the cell can perform its functions, corresponding to ‘neural cell classes’. Here we described such a grouping of neurons, as we identify four putative inhibitory and six putative excitatory cell classes in the cortical layers 2/3 of adult S1 and a similar grouping in V1, with 7 putative inhibitory sub-classes, and 4 putative excitatory sub-classes.

Neurons can plausibly adopt different strategies in order to perform certain functions (37), resulting in gradual differences between neurons rather than well-separated ‘clusters’ of properties as we observed among the excitatory neurons (37). Finding the neural properties that group together, i.e. ‘clustering’ our high-dimensional dataset, is a non-trivial task, where the experimenter necessarily has to make non-obvious choices about which clustering method to use and how to set the meta-parameters of the clustering algorithm. Often, the ‘ground-truth’ is not known, so it is difficult to assess the quality of the chosen method. Therefore, it is important to choose the method with some knowledge of the dataset (39). For instance, because of the continuous nature of the data examined, clustering methods purely based on distance, such as tSNE (40), do not perform well. An extra difficulty in our datasets is that they are large for ex vivo datasets, but relatively small in relation to the number of dimensions. Therefore, we chose a combination of a simple but well-understood and easy-to-interpret automatic method (k-means clustering) together with a method that was specific to this dataset (dismembering small clusters and re-assigning each element separately until each cluster had at least 10 neurons). This approach allowed us to perform clustering in a lower dimensional space, i.e. by considering a lower number of features. Repeating the clustering in a larger dataset, if it becomes available, will help to identify those neuronal clusters that were sparsely populated in our analysis.

While generally, we found a similar classification between the somatosensory and visual cortex datasets, there are some differences between the two. The most obvious one, i.e. the higher number of clusters found in the V1 dataset, might be attributed to the arbitrary choice of the minimum number of cells per cluster that we took and an extra cluster might have passed that threshold, possibly also thanks to the slightly higher number of cells included in the V1 dataset. The higher number of descriptive variables that survived the correlation-based dimensionality reduction might instead be due both to an intrinsically greater diversity in the neuronal response, and to the higher noise level present in the experiments from the visual cortex database with respect to the somatosensory one.

Even though the high-dimensional clustering of cell properties gives many insights on its own, it is not always feasible in an experimental setup to measure so many cellular properties. Therefore, we asked: ‘How can we minimize the number of measured cell properties, and still perform a cluster analysis with the same results as the high-dimensional clustering?’. For this, we had to perform dimensionality reduction. With the recent improvement of high-dimensional recording techniques, dimensionality reduction methods beyond ‘simple’ principal component analysis have moved into the spotlight. Even though these types of analysis are mostly used for population recordings (41; 42), the same steps apply for high-dimensional in vitro single-cell data. The first step is looking into the ‘underlying’ dimensionality of the dataset: one should check that even though a (typically large) dataset is high-dimensional, the underlying principles are probably low-dimensional, allowing for a low-dimensional description. We showed here through the dimensionality calculation and the correlations between the measured parameters in our dataset that a lower-dimensional representation should be possible (Figure 1). The second step is to identify informative and less informative dimensions (Figure 2). Finally, one should perform a check on the dimensionality reduction: does the representation in low-dimensional space yield the same properties as in the high-dimensional space? Here we performed our clustering analysis both on the high- and low-dimensional representations of our data, and showed that neurons are classified into well-separated clusters before and after the dimensionality reduction. Thus, the clustering approach presented herein is unlikely to be biased by the sampling size or the number of features used for the classification. Future application of this clustering approach in neurons recorded across brain regions and developmental periods will help to address whether these cell classes are universal.

The fact that the performance of each feature in determining each cluster was strongly dependent on the stimulus intensity, should first of all act as a reminder of the necessity to contextualize the results of a classification effort, and of the growing need for standardization of experimental protocols. Further, this acts as a confirmation of the importance to include dynamical parameters as descriptors of neuronal activity. The most interesting part of the result might however be how it strongly suggests that the functional properties of neuronal sub-types, and therefore their activity, are dependent on stimulus properties.

Heterogeneity is hypothesized to improve coding (48; 49; 50). Indeed, it is known that different groups of neurons contribute differently to the representation of external stimuli with varying properties such as firing sparseness or temporally precise responses (43; 44): a functional clustering based on the integrative and firing properties, such as the one we propose, would help in identifying these neuronal groups, characterize them, and possibly provide a link between their stimulus representation and their biophysical properties. Using the identified cell classes presented here and their response properties in computational network models will help to develop more advanced, biologically realistic networks, starting with a model of the somatosensory cortex of the rodents, to match the detailed V1 already presented by the Allen Institute (47).

## Acknowledgments

This work was supported by grants from the European Commission (Horizon2020, nr. 660328), European Regional Development Fund (MIND, nr. 122035) and the Netherlands Organisation for Scientific Research (NWO-ALW Open Competition, nr. 824.14.022) to TC, the Netherlands Organisation for Scientific Research (NWO-Veni, nr. 863.150.25) to FZ, and a doctoral fellowship by the National Council for Scientific and Technological Development of Brazil (CNPQ) to ASL.

## Appendix 1

### Supplementary Figures

**Appendix 1—figure 1.**
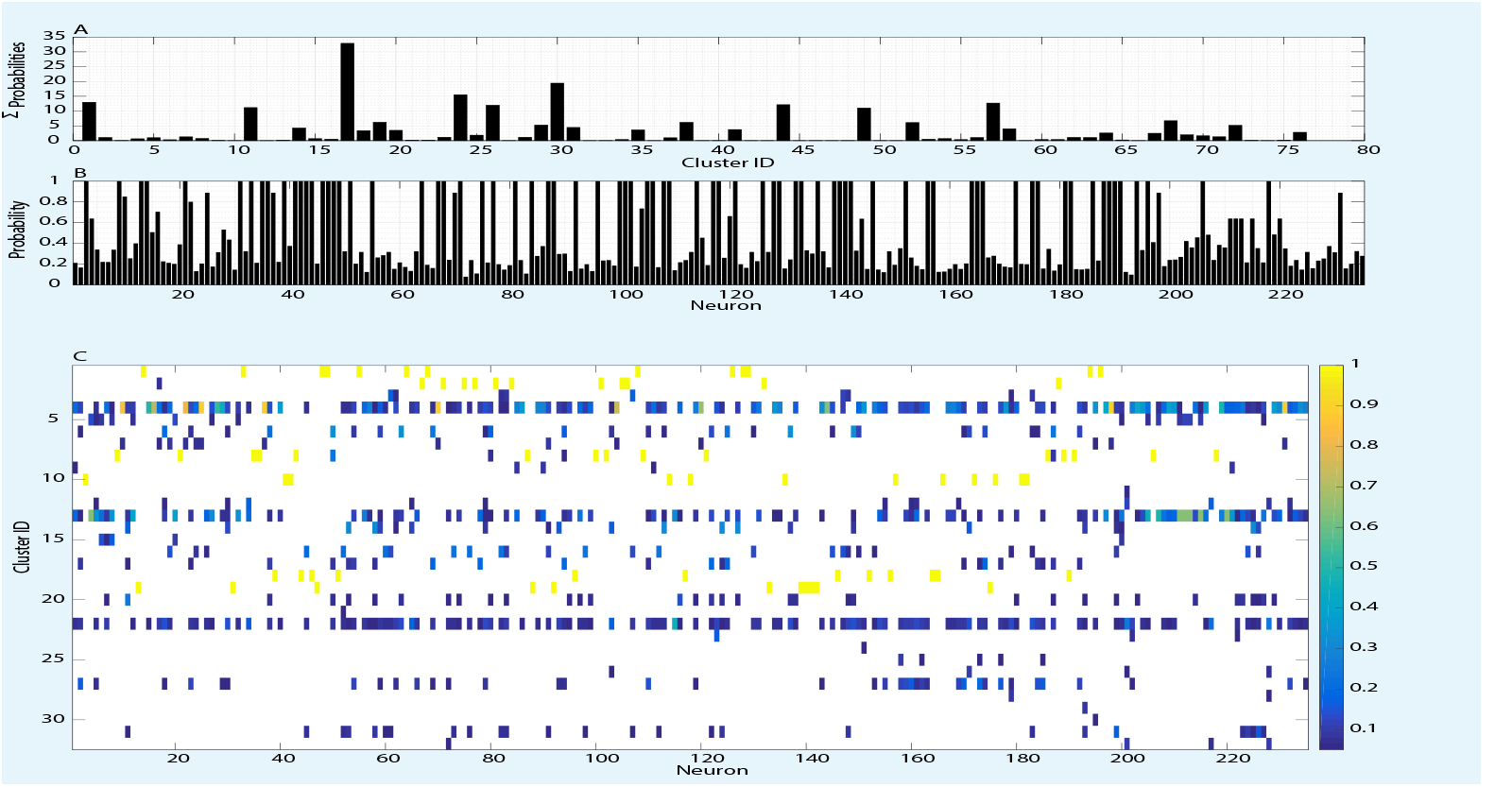
Clustering in high-dimensional feature space is a probabilistic approach to cell-type identification. **(A)** Sum of the probabilities of all the neurons to belong to a specific cluster. **(B)** The maximum probability for each neuron to belong to any cluster (indicative of certainty of cluster assignment). **(C)** The probability of each cell to be assigned to a specific cluster after the merging procedure, counted after 10000 mergings iterations.

**Appendix 1—figure 2.**
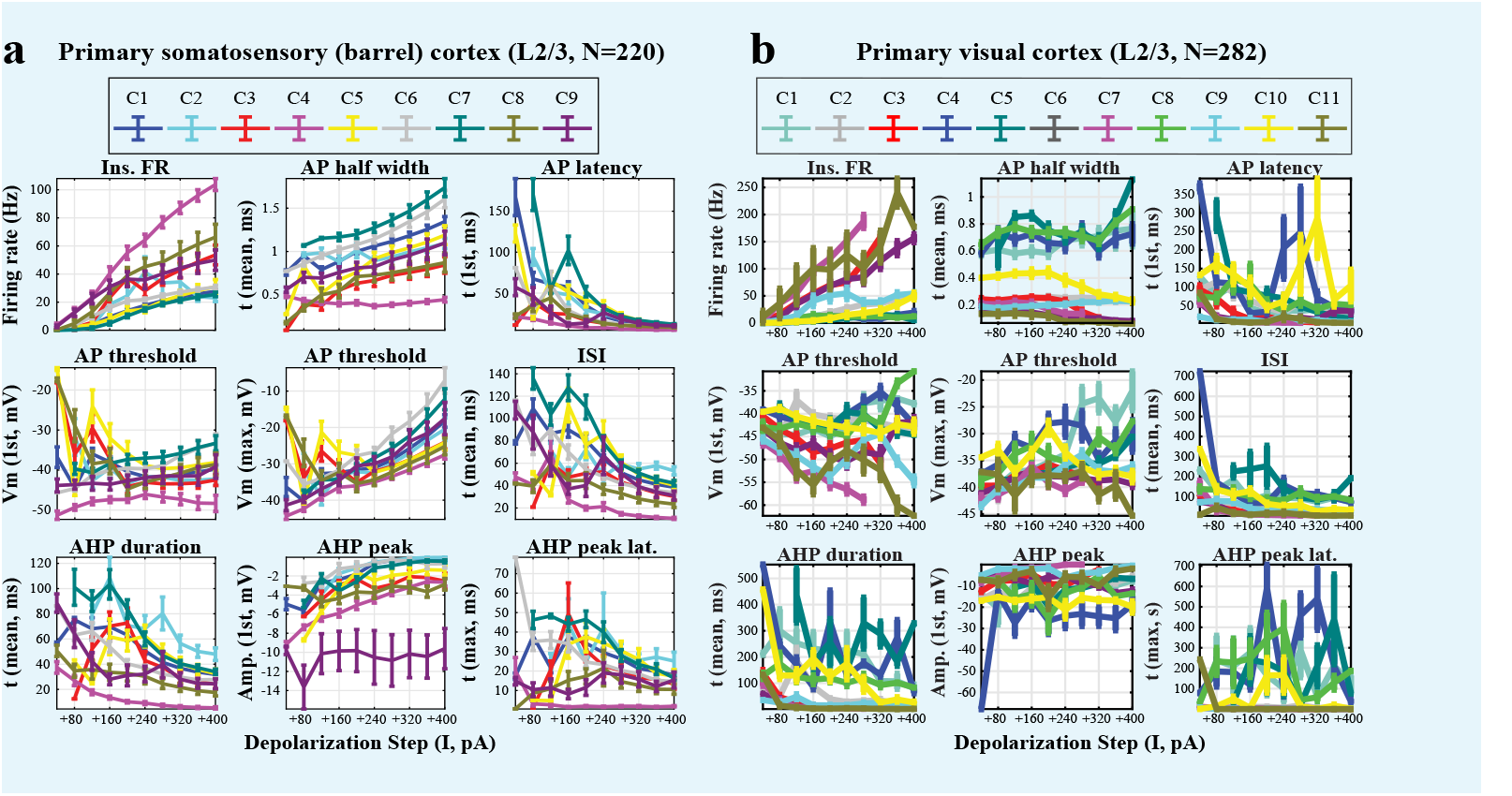
Characteristics of the identified clusters in the primary somatosensory (barrel) and visual cortices of mouse. The clusters were obtained via k-means on the subspace of the 5 parameters found to be most descriptive of each datasets.

### Supplementary Tables

**Appendix 1—table 1.** Principal features of action potentials. Each quantity is measured over increasing current injection steps. For each quantity (except the 1st, which has only a single scalar value), both the starting value and the slope of the linear fit over the current injections are used as parameters (for a total of 139 parameters). AP = action potential; AHP = afterhyperpolarization current; ISI = interspike interval. The 40 and 48 parameters that were selected after the correlation analysis in the S1 and V1 databases are in bold and italics, respectively.

| Feature | Description |
| --- | --- |
| 1. Current necessary for 1st AP [pA] | Lowest current step at which the cell fired an AP |
| 2. AP count | Number of APs fired through the stimulation |
| 3. Absolute FR [Hz] | Number of APs divided by the stimulus duration |
| 4. Instantaneous FR [Hz] | Instantaneous firing rate over the stimulation |
| 5. AP Latency of 1st spike [ms] | Time between the start of stimulation and the 1st AP |
| 6. AP Latency of last spike [ms] | Time between the start of stimulation and the last AP |
| 7. AP Latency window [ms] | Interval between the first and last AP generated |
| 8. AP Latency compression | Onset latency between the first and last spikes, divided by the stimulus length |
| 9. ISI minimum [ms] | Minimum ISI between two consecutive APs over the stimulation |
| 10. ISI maximum [ms] | Maximum ISI between two consecutive APs over the stimulation |
| 11. ISI mean [ms] | Mean ISI between two consecutive APs over the stimulation |
| 12. ISI median [ms] | Median ISI between two consecutive APs over the stimulation |
| 13. ISI adaptation rate [%] | (First ISI - Last ISI) / First ISI |
| 14. AP Threshold of 1st spike [mV] | Threshold at which the 1st AP is generated, computed as the peak of the 2nd derivative of the membrane potential. |
| 15. AP Threshold of last spike [mV] | Threshold at which the last AP is generated, computed as the peak of the 2nd derivative of the membrane potential. |
| 16. AP Threshold minimum [mV] | Minimum threshold at which an AP is generated during the AP train. |
| 17. AP Threshold maximum [mV] | Maximum threshold at which an AP is generated during the AP train. |
| 18. AP Threshold mean [mV] | Mean threshold at which an AP is generated during the AP train. |
| 19. AP Threshold median [mV] | Median threshold at which an AP is generated during the AP train. |
| 20. AP Threshold adaptation rate [%] | (First AP Threshold - Last AP Threshold) / First AP Threshold |
| 21. AP Half-Width of 1st spike [ms] | Width of the 1st AP at half height |
| 22. AP Half-Width of last spike [ms] | Width of the last AP at half height |
| 23. AP Half-Width minimum [ms] | Minimum half-width of the whole AP train |
| 24. AP Half-Width maximum [ms] | Maximum half-width of the whole AP train |
| 25. AP Half-Width mean [ms] | Mean half-width of the whole AP train |
| 26. AP Half-Width median [ms] | Median half-width of the whole AP train |
| 27. AP Half-Width adaptation rate [%] | $(\text{First AP Half-Width} - \text{Last AP Half-Width}) / \text{First AP Half-Width}$ |
| 28. AP Amplitude of 1st spike [mV] | Amplitude of the 1st AP, computed as $\vert \text{max} - \text{baseline}\vert $ |
| 29. AP Amplitude of last spike [mV] | Amplitude of the last AP, computed as $\vert \text{max} - \text{baseline}\vert $ |
| 30. AP Amplitude minimum [mV] | Minimum amplitude of the whole AP train |
| 31. AP Amplitude maximum [mV] | Maximum amplitude of the whole AP train |
| 32. AP Amplitude mean [mV] | Mean amplitude of the whole AP train |
| 33. AP Amplitude median [mV] | Median amplitude of the whole AP train |
| 34. AP Amplitude adaptation rate [%] | $(\text{First AP Amplitude} - \text{Last AP Amplitude}) / \text{First AP Amplitude}$ |
| 35. AHP Peak Latency of 1st spike [ms] | Time interval between the return to baseline of the voltage after the 1st AP of the train and its corresponding AHP peak |
| 36. AHP Peak Latency of last spike [ms] | Time interval between the return to baseline of the voltage after the last AP and its corresponding AHP peak |
| 37. AHP Peak Latency minimum [ms] | Minimum time interval between the return to baseline and the AHP peak |
| 38. AHP Peak Latency maximum [ms] | Maximum time interval between the return to baseline and the AHP peak |
| 39. AHP Peak Latency mean [ms] | Mean time interval between the return to baseline and the AHP peak |
| 40. AHP Peak Latency adaptation rate [%] | $(\text{First AHP Peak Latency} - \text{Last AHP Peak Latency}) / \text{First AHP Peak Latency}$ |
| 41. AHP Peak Amplitude of 1st spike [mV] | Difference between the peak of the AHP corresponding to the 1st AP and the baseline membrane potential |
| 42. AHP Peak Amplitude of last spike [mV] | Difference between the peak of the AHP corresponding to the last AP and the baseline membrane potential |
| 43. AHP Peak Amplitude minimum [mV] | Minimum difference between the peak of the AHP corresponding to the APs and the baseline membrane potential |
| 44. AHP Peak Amplitude maximum [mV] | Maximum difference between the peak of the AHP corresponding to the APs and the baseline membrane potential |
| 45. AHP Peak Amplitude mean [mV] | Mean difference between the peak of the AHP corresponding to the APs and the baseline membrane potential |
| 46. AHP Peak Amplitude adaptation rate [%] | $(\text{First AHP Amplitude} - \text{Last AHP Amplitude}) / \text{First AHP Amplitude}$ |
| 47. AHP Duration of 1st spike [ms] | Time interval between the membrane potential reaching baseline after an AP and reaching baseline again after the corresponding AHP, measured for the 1st AP |
| 48. AHP Duration of last spike [ms] | Time interval between the membrane potential reaching baseline after an AP and reaching baseline again after the corresponding AHP, measured for the last AP |
| 49. AHP Duration minimum [ms] | Minimum time interval between the membrane potential reaching baseline after an AP and after the corresponding AHP |
| 50. AHP Duration maximum [ms] | Maximum time interval between the membrane potential reaching baseline after an AP and after the corresponding AHP |
| 51. AHP Duration mean [ms] | Mean time interval between the membrane potential reaching baseline after an AP and after the corresponding AHP |
| 52. AHP Duration adaptation rate [%] | $(\text{First AHP Duration} - \text{Last AHP Duration}) / \text{First AHP Duration}$ |
| 53. AHP Integral of 1st spike [ms*mV] | Integral of the AHP of the 1st AP in the train |
| 54. AHP Integral of last spike [ms*mV] | Integral of the AHP of the last AP in the train |
| 55. AHP Integral minimum [ms*mV] | Minimum integral of the AHP over the train |
| 56. AHP Integral maximum [ms*mV] | Maximum integral of the AHP over the train |
| 57. AHP Integral mean [ms*mV] | Mean integral of the AHP over the train |
| 58. AHP Integral adaptation rate [%] | $(\text{First AHP Integral} - \text{Last AHP Integral}) / \text{First AHP Integral}$ |
| 59. AHP Fall Slope of 1st spike [mV/ms] | Slope of the decay phase of the AHP of the 1st AP in the train |
| 60. AHP Fall Slope of last spike [mV/ms] | Slope of the decay phase of the AHP of the last AP in the train |
| 61. AHP Fall Slope minimum [mV/ms] | Minimum slope of the decay phase of the AHP in the train |
| 62. AHP Fall Slope maximum [mV/ms] | Maximum slope of the decay phase of the AHP in the train |
| 63. AHP Fall Slope mean [mV/ms] | Mean slope of the decay phase of the AHP in the train |
| 64. AHP Fall Slope adaptation rate [%] | $(\text{First AHP Fall Slope} - \text{Last AHP Fall Slope}) / \text{First AHP Fall Slope}$ |
| 65. AHP Rise Slope of 1st spike [mV/ms] | Slope of the rising phase of the AHP of the 1st AP in the train |
| 66. AHP Rise Slope of last spike [mV/ms] | Slope of the rising phase of the AHP of the last AP in the train |
| 67. AHP Rise Slope minimum [mV/ms] | Minimum slope of the rising phase of the AHP in the train |
| 68. AHP Rise Slope maximum [mV/ms] | Maximum slope of the rising phase of the AHP in the train |
| 69. AHP Rise Slope mean [mV/ms] | Mean slope of the rising phase of the AHP in the train |
| 70. AHP Rise Slope adaptation rate [%] | $(\text{First AHP Rise Slope} - \text{Last AHP Rise Slope}) / \text{First AHP Rise Slope}$ |

